# The age and mouse sperm quality – a flow cytometry investigation

**DOI:** 10.1101/2021.07.18.451583

**Authors:** Federica Zacchini, Michal Bochenek, Simona Bisogno, Alan Chan, Grazyna Ewa Ptak

**Affiliations:** Malopolska Centre for Biotechnology, Jagiellonian University, Krakow, Poland; Percuros BV, Leiden, The Netherlands

**Keywords:** Aging, Spermatozoa, DNA, Mouse

## Abstract

Postponement of fatherhood is growing worldwide due to socio-economic factors. The choice to conceive the first child above the age of 35 years is often associated with reduced fertility and poor pregnancy outcome. As widely known, several factors (*e.g.*, lifestyle, environment, health problems) can affect spermatogenesis leading to poor reproductive outcome. Currently, the debate on the influence of aging on male gametes and safety/risk of conception at advanced age is still ongoing. Controversial results have been published so far on the changes in semen features of aging men and other mammalian species (mainly rodents). In this study, we aimed to assess how aging affects sperm quality in an inbreed mouse model, without underlying infertility, using a flow cytometry approach. Our data showed that aging is associated with increased sperm chromatin condensation, but not changes in the DNA integrity, metabolic activity or viability. These data suggest a mild effect of aging on sperm quality in a mouse model without underlying infertility.

## 1. Introduction

In the last decades, social and economic factors as well as the increased life expectancy and the development of assisted reproductive techniques contribute to the postponement of parenthood. A growing number of studies suggested conception at advanced paternal age (APA) as risk factor for increased time to pregnancy, increased miscarriage rate and, in the long term, predisposition to adult diseases in the offspring [1–3]. In human, the observed poor pregnancy outcome due to APA has been associated with decreased sperm quality (*e.g.*, low sperm concentration and motility, high DNA fragmentation) [4,5]. However, data from human studies cannot fully discern the effect of aging from that of concomitant confounding factors (*i.e.*, obesity, diabetes, smoking, exposure to toxins and environmental pollution). Also, a systematic and comparative evaluation of the outcome of different studies is difficult due to the lack of an optimal definition of APA, resulting in different age cut-off among studies. Similarly, in the last decade animal studies provided various results, which make difficult to determine how and to what extent age affects sperm quality and functions. For example, Kotarska et al,2017 [6] described a correlation between age and reduced sperm quality and reproductive fitness in subfertile B10.BR-Y^del^ mice while no differences were observed between young vs old wild type B10.BR mice. Similarly, strong effect of aging on semen parameters has been described in mutant mice but only mild effect in wildtype ones [7]. In our previous studies, we observed that APA was not associated with reduced fertility, in terms of pregnancy rate obtained after mating of old males with young females, while behavioural abnormalities were observed in resulting offspring [8]. Other study described reduced fertility in mice > 12 months old but not changes in qualitative sperm parameters (*e.g.*, concentration, motility) [9]. One explanation can be the occurrence of changes (*i.e*., damages, mutations) in the sperm DNA rather than qualitative parameters (*e.g*., viability, motility or concentration), which are compatible with *in utero* life but may interfere with developmental programming and contribute to increase incidence of post-natal diseases. In the current study, our aim was to assess the influence of aging on spermatozoa in a mouse model, without underlying infertility, using a flow cytometry approach. Inbreed C57BL6 fertile males at age ranging from 2 months (corresponding to 20 years in human) to 15 months (corresponding to > 60 years in human) were used. Individual sperm samples were analysed by flow cytometry for the following parameters: i) DNA fragmentation, breaks (both single and double strand) and chromatin condensation, ii) metabolic activity and iii) apoptosis. Our data overall indicate a mild effect of aging on sperm quality in a mouse model without underlying infertility.

## 2. Material and methods

### 2.1 Ethical statement and animal maintenance

Animal breeding and experimental procedures were conducted according to EU directive 86/609 and to national law for Animal Experimentation. C57BL6 mice were maintained at the animal facilities of the Institute of Zoology of Jagiellonian University in Cracow. Animals were housed in conventional cages with water and food ad libitum in a room with 12h dark/light cycle, 20 ± 2 °C and humidity >50%. Animals were housed in groups (maximum 5 animals/cages) and environmental enrichment provided.

### 2.2. Collection of spermatozoa from cauda epididymis

Sperm were isolated from 216 inbreed C57BL6 mice at age between 2 and 15 months, as previously described [10] with minor modification. Briefly, caudae epididymis were isolated from each male and transferred into 500 μl of G-MOP Plus (Vitrolife) at 37ºC. Spermatozoa were released by squeezing and puncturing the cauda epididymis using a needle. Then, spermatozoa were allowed to swim out for 20 min at 37ºC. Individual sperm samples (mean 47±15 × 10^6^ sperm/ml) were divided into 3 parts used for all 3 analyses described below, such as Sperm Chromatin Structure Assay (SCSA), CellROX Green assay (Invitrogen) for metabolic activity and AnnexinV/FITC assay (Beckman Coulter) for apoptosis.

To avoid bias within the study: i) the animal have been maintained under controlled conditions (as described in section 2.1), ii) in case of health problems, animals have been removed from the study, iii) individual samples have been subjected to all analysis described below, iv) at least three replicates (with at least 3 samples/replicate) have been carried out for each analysis.

### 2.3 Sperm DNA fragmentation and chromatin status

The modified Sperm Chromatin Structure Assay (SCSA) method was used to examine sperm DNA fragmentation and chromatin condensation, as previously described [11]. Briefly, individual sperm samples (5 × 10^6^ cells/ml) were incubated with HCl solution (pH=1.5) for 30 seconds on ice. Then, samples were stained with acridine orange for 3 minutes on ice. The green (normal DNA) and red (fragmented DNA) fluorescence signals were collected from 12 000 spermatozoa. A ‘Sperm DNA Fragmentation’ ratio parameter was created according to formula: red/(green+red) fluorescence. This ratio parameter allows to present normal DNA and even very low level of DNA fragmentation. The high and moderate DNA Fragmentation Index (DFI) was calculated from Sperm DNA Fragmentation histogram (Figure 1E). Also, poor chromatin condensation was assessed from green vs sperm DNA fragmentation dot plot (Figure 1F) as spermatozoa with higher acridine orange uptake. In addition to original SCSA data processing, a novel gating strategy was applied to green vs. sperm DNA fragmentation dot plot. The population of spermatozoa with fragmented DNA was divided into 2 groups: a) with normal green fluorescence and expected dominance of DNA single strand breaks (ssbDNA) as at lower level of DNA denaturation and b) low green fluorescence thus with expected double strand breaks (dsbDNA) - higher level of DNA denaturation (Figure 1I). Fluorescence measurements were carried out using Navios flow cytometer (Beckman Coulter), as described in section 2.6.

**Figure 1:**
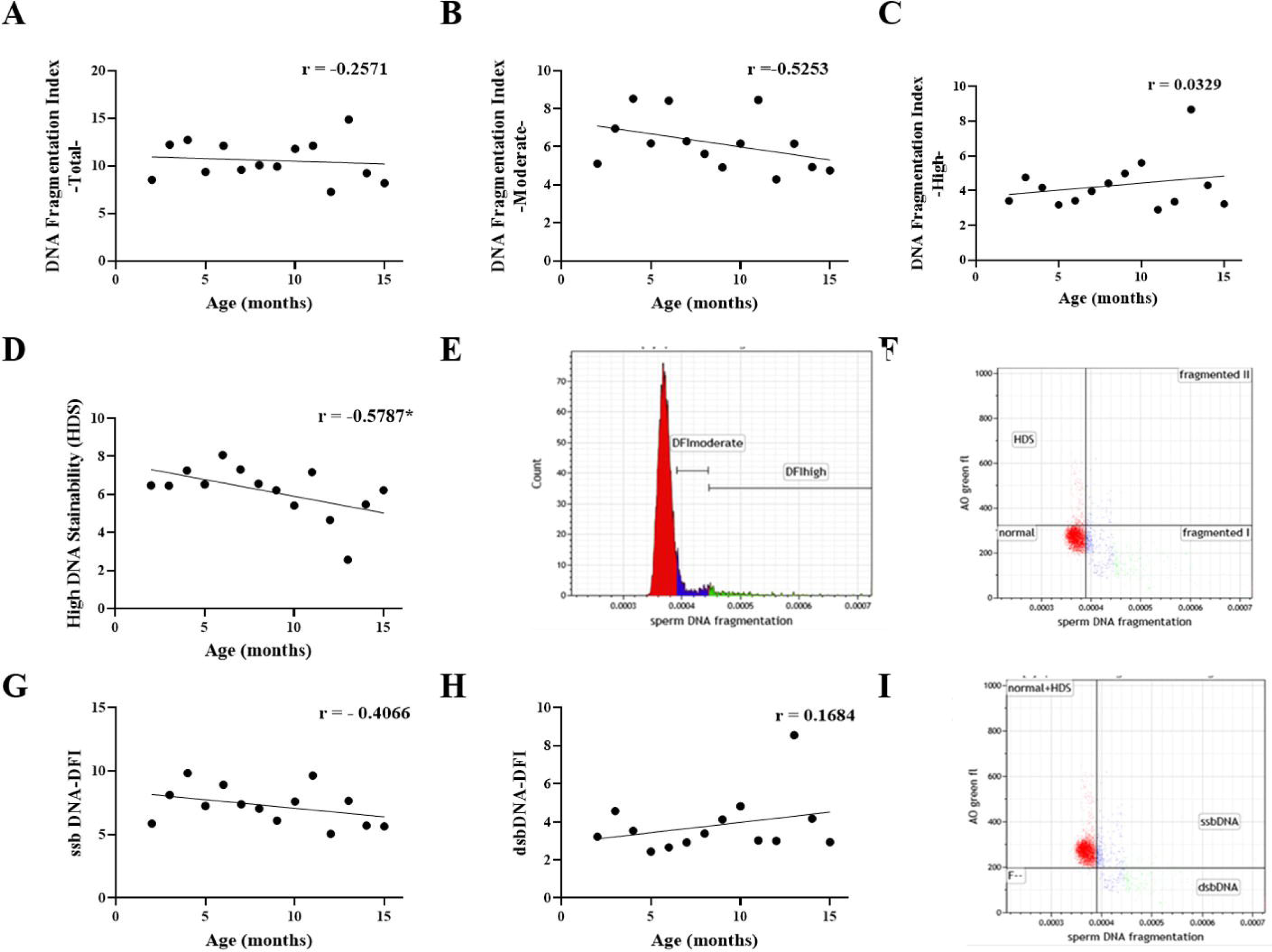
Increasing DNA condensation but not DNA fragmentation is associated with aging in mouse spermatozoa. Sperm Chromatin Structure Assay (SCSA) showed that aging was not significantly correlated neither with the total level of DNA fragmentation (A) not with moderate (B) and high (C) DNA fragmentation in spermatozoa obtained from mouse at age between 2 and 15 months. (D) Significant negative correlation between High DNA Stainability (HDS) and mouse age (Spearman r= − 0.5787, * is P = 0.032), indicative of increased DNA condensation in aged mice. (E) SCSA cytogram example for determination of DNA Fragmentation Index (DFI). (F) SCSA cytogram example for determination of HDS. (G-H) Aging was not associated with single strand DNA breaks (ssbDNA) while a weak (r = 0.1684), but not statistically significantly (P = 0.057) correlation, between aging and DNA double strand breaks (dsbDNA) and was found. (I) Cytogram example for the assessment of ssbDNA. n > 13 samples at age 2-12 and 15 months, n ≥ 3 samples at age 13 and 14.

### 2.4 Metabolic activity assay

To assess the influence of age on metabolic activity expressed as intracellular ROS level of mouse sperm, CellROX Green assay (Invitrogen) has been used according to manufacturer’s instructions, with minor modification. Briefly, sperm fraction was diluted 1:10 in Phosphate Buffered Saline (PBS, Lonza) and CellROX reagents (1 μM/μl) added. The solution was incubated at 37 °C for 30 minutes. Then, 2 μl of 2.4 mM propidium iodide (PI) solution were added. After 2 minutes of incubation at room temperature, the green (for CellROXgreen) and red (for PI) fluorescence were collected from 12 000 spermatozoa. Fluorescence measurements were carried out using Navios flow cytometer (Beckman Coulter), as described in section 2.6. The sperm population was divided into 6 subpopulations: i) live and active, ii) moribund and active, iii) dead and active, iv) live and inactive, v) moribund and inactive, vi) dead and inactive.

### 2.5 Apopotosis Assay

AnnexinV/FITC Kit (Beckman Coulter) was used to quantify apoptosis in mouse sperm, according to manufacturer’s instructions, with minor modification. Briefly, AnnexinV reagent was added to sperm fraction diluted 1:10 in PBS (LONZA) and the solution incubated on ice for 15 minutes. Then, 400 μl of buffer with 2 μl of 2.4 mM PI solution (instead of the Kit’s 7AAD fluorochrome) was added. After 2 minutes of incubation at room temperature, the green (for Annexin V) and red (for PI) fluorescences were collected from 12 000 spermatozoa. Fluorescence measurements were carried out using Navios flow cytometer (Beckman Coulter), as described in section 2.6. Individual sperm populations were divided into the following subpopulations: i) live spermatozoa, ii) apoptotic spermatozoa and iii) necrotic spermatozoa

### 2.6 Flow cytometry measurement

For all assays in paragraph 2.3, 2.4 and 2.5, the Navios (Beckman Coulter) flow cytometer was used in the following configuration: the blue, 488nm laser for fluorochromes excitation; filter set of 550DCSP/525BP for green fluorescence and 655DCSP/620BP for red fluorescence. The ‘Medium’ sheath flow rate was applied for all samples and 12 000 spermatozoa were acquired. Instrument calibration was checked each day with SPHERO Ultra Rainbow (Spherotech) calibration particles. For off-line data analyses the Kaluza 2.1 (Beckman Coulter) software was used. For SCSA, to exclude non-cellular debris the following 2 gates were applied: on Forward Scatter vs Side Scatter (both logarithmic) and on green vs red fluorescence (both linear) cytograms. Additionally, the third gate on red fluorescence vs time cytogram was applied (if flow disturbances were recognized during sample acquisition). No fluorescence compensation was used between green and red channels. For CellROX assay, to exclude non-cellular debris the gate on Forward Scatter vs Side Scatter (both logarithmic) was used. The red (PI) channel was compensated by 2.1%. For AnnexinV assay, to exclude non-cellular debris the gate on Forward Scatter vs Side Scatter (both logarithmic) was used. The red (PI) channel was compensated by 0.7%.

### 2.7 Data analysis

Raw data were analysed using GraphPad Prism 8. Pearson’s correlation was used to assess the influence of age on sperm parameters. Values of P < 0.05 were considered statistically significant.

## 3 Results

### 3.1 Increased DNA condensation but not DNA fragmentation in aging mouse spermatozoa

Figure 1 shows that aging in mouse does not affect the total DNA fragmentation (Fig. 1A) as well as the moderate and high DNA fragmentation levels (Fig. 1B-C). Figure 1D shows a significant reduction of High DNA Stainability (HDS) over time (r = − 0.5787, P = 0.03), indicative of increasing chromatin condensation. Then, by using a novel gating strategy, the level of DNA single strand and double strand breaks (ssbDNA and dsbDNA, respectively) were assessed. No correlation was observed between aging and ssbDNA (Fig. 1G) while a weak (r = 0.1684) but not statistically significant (P > 0.05) correlation was found between aging and dsbDNA (Fig. 1H) in mouse sperm.

### 3.2 Aging is not associated with impaired metabolic activity in mouse spermatozoa

Metabolic activity was evaluated in mouse spermatozoa using CellROX Green assay. The flow cytometry analysis was performed on the same samples used for SCSA. Figure 2 shows a moderate negative, but not statistically significant- correlation between age and the percentage of moribund/dead active subpopulation (r = − 0.4251 and − 0.5127, respectively; P = 0.06, Fig 2B-C) while no effect of aging on the distribution of the other subpopulations (Fig2A-B, D-F) in mouse spermatozoa.

**Figure 2:**
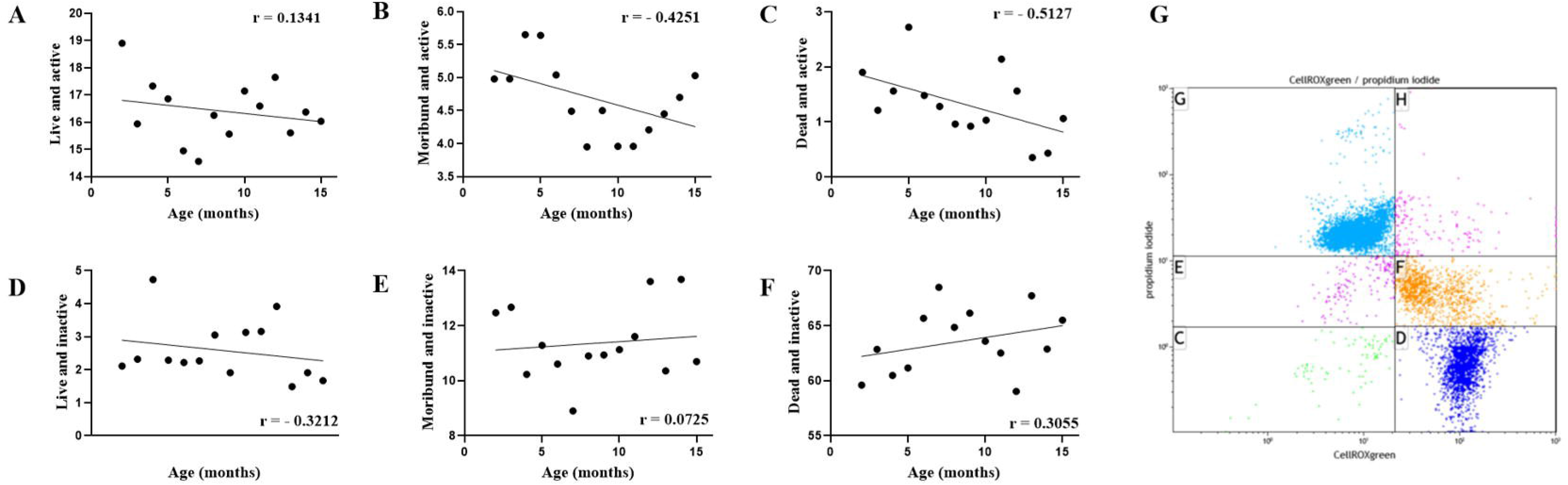
Aging does not affect metabolic activity of mouse spermatozoa. CellROX Green assay did not reveal significant influence of aging on the metabolic activity of mouse spermatozoa. (A) Graph shows comparable level of live and active spermatozoa among groups. (B) Graph shows moderate negative but not statistically significant (P > 0.05) correlation between level of moribund-active spermatozoa and aging. (C) Graphs shows a negative, but not statistically significant, correlation between the rate of dead-active spermatozoa associated with aging (r = −0.5127, P = 0.06) (D) Graph shows comparable level of live and inactive spermatozoa among groups. (E) Graph shows comparable level of moribund-inactive spermatozoa among groups. (F) Graph shows comparable level of dead inactive spermatozoa among groups. (G) Cytogram of metabolic activity (that is intracellular ROS activity) and membrane integrity assay (CellROX Green vs propidium iodide). Regions description: C – live, ROS inactive; D – intact membranes, ROS active; E – moribund, ROS inactive; F – moribund, ROS active; G – dead, ROS inactive; H – damaged membranes, ROS active n > 13 samples at age 2-12 and 15 months, n ≥ 3 samples at age 13,14.

### 3.3 Apoptosis is not induced by aging in mouse spermatozoa

To assess whether aging lead to increased apoptotic cell death in mouse, AnnexinV/FITC + Propidium Iodide staining has been carried out on the same samples used for SCSA and metabolic activity. Figure 3A-B shows no effect of aging in the percentage of live and apoptotic spermatozoa in our mouse model. Figure 3C shows a weak but not statistically significant increase of necrotic subpopulation in mouse spermatozoa over time (r = 0.2359, P > 0.05).

**Figure 3:**
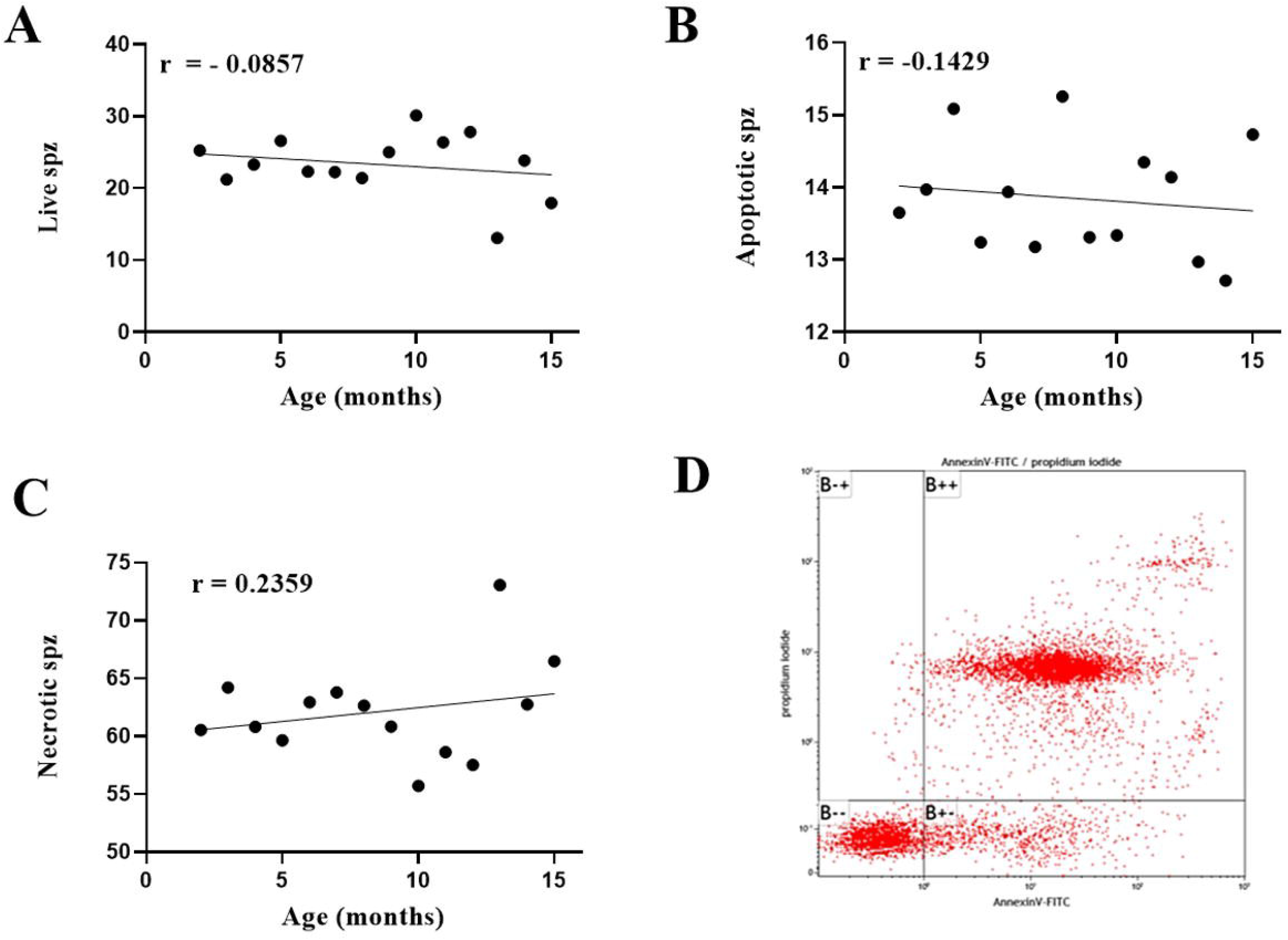
Aging does not induce apoptotic cell death in mouse spermatozoa. AnnexinV/FITC assay of mouse spermatozoa from 2 to 15 months of age showed comparable level of apoptosis among groups. (A) Graph shows percentage of live spermatozoa. Please note a weak negative – not statistically significant-correlation between mouse age and percentage of live spermatozoa (r = −0,2571, P > 0.05). (B) Graph shows comparable level of apoptosis among groups. (C) Graph shows a moderate – but not statistically significant - positive correlation between age and necrotic spermatozoa (r = 0.2359, P = 0.4). (D) Cytogram for the Apoptosis (phosphatidylserine externalization) assay (AnnexinV/FITC vs propidium iodide). Regions description: B−−: live; B+−: apoptotic; B−+ and B++: necrotic. n > 13 samples at age 2-12 and 15 months, n ≥ 3 samples at age 13,14.

### 3.4 Influence of temperature on membrane integrity

Membrane integrity in aging spermatozoa has been assessed by staining with Propidium Iodide within CellROX assay, after incubation at 37□/30 minutes, and independently within Annexin V/FITC assay, following incubation on ice/15 minutes. The same concentration of PI (2 μl of 2.4 mM solution) was used and 12 000 spermatozoa analyzed within the two assays. The two measurements of membrane integrity did not overlap for individual samples and show different values (Figure 4), suggesting high susceptibility to environmental conditions.

**Figure 4:**
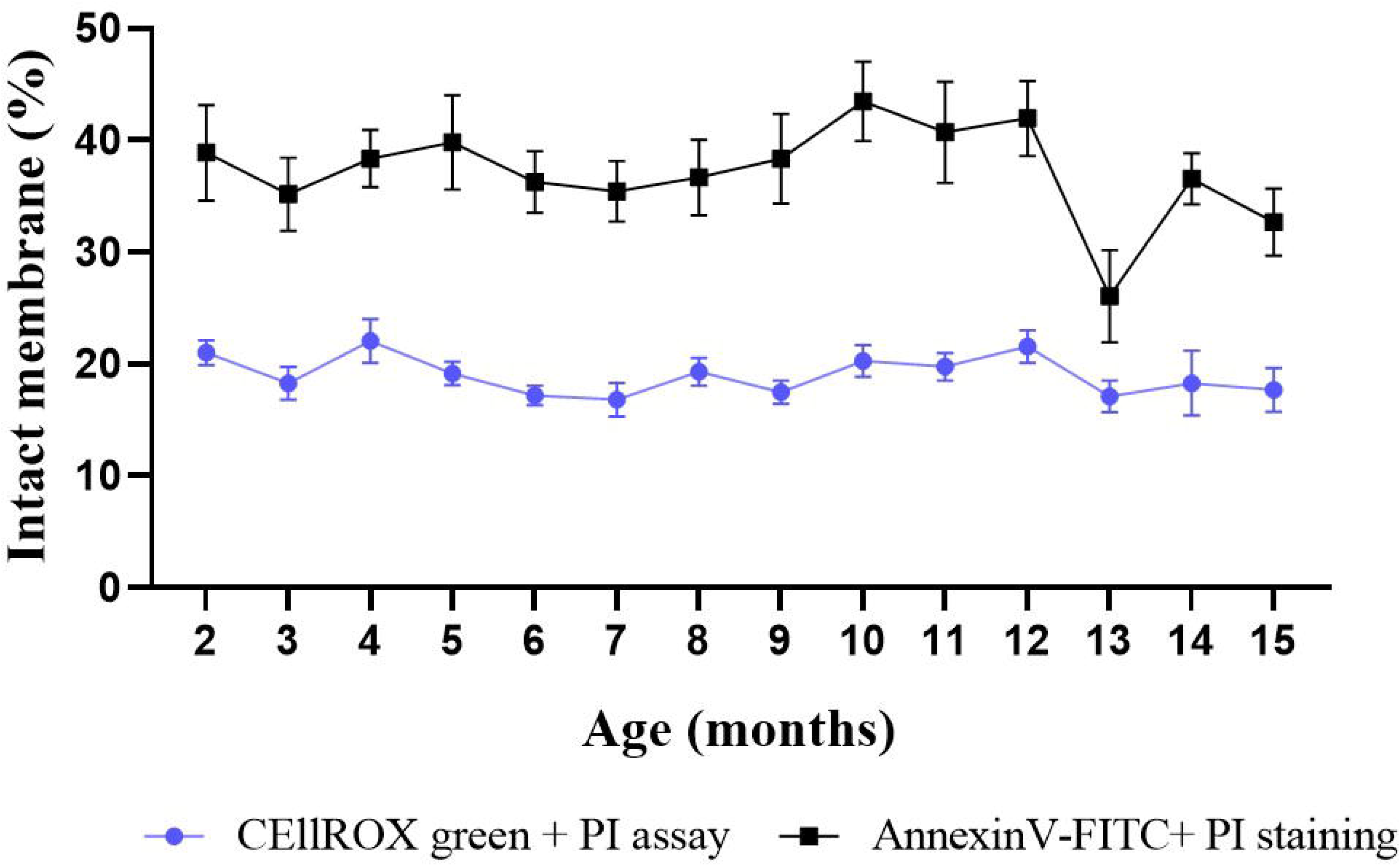
Changes in membrane integrity due to temperature in mouse spermatozoa. Individual mouse sperm samples were stained with propidium iodide, marker of membrane integrity, after incubation 30 minutes at 37□ (for CellROX green assay) or 15 minutes on ice (for AnnexinV-FITC staining). Graphs shows that the measurements of membrane integrity did not overlap between the CellROX green+PI assay and Annexin V-FITC+PI staining, suggesting high susceptibility to environmental conditions. n > 13 samples at age 2-12 and 15 months, n ≥ 3 samples at age 13,14 months.

## 4 Discussion

In the present study, the influence of aging on sperm quality in a mouse model without underlying infertility was evaluated using a flow cytometry approach. Our findings showed that aging is associated with increased DNA condensation but not with DNA fragmentation or perturbation of metabolic activity and viability in mouse spermatozoa from inbreed C57BL6 mice.

It is widely described that sperm DNA integrity is a crucial factor influencing proper embryo/fetal development and, in the long term, offspring health. Previous studies described poor embryo development [12–14] and increased risk of adult diseases (mainly neurodevelopmental disorders) in APA offspring [8, 15–17]. It has been suggested that the above-mentioned side effects of APA might be associated with abnormalities occurring in sperm DNA (*e.g.*, de novo mutation, epigenetic defects) [12, 15, 18]. Notwithstanding, we did not observed changes in DNA fragmentation, our data showed that aging was associated with increased chromatin condensation over time in mouse sperm. One of the mechanisms behind the observed high condensation may be the increasing methylation in aging sperm, recently described by us and others [15, 19]. The occurrence of epigenome changes in aging sperm is of great concern. In fact, a growing number of studies reported that epigenetic defects are risk factor for the onset of post-natal diseases (*i.e.*, neurodevelopmental disorders, metabolic syndrome) in the offspring, which may be transmitted across generations [15, 20–22]. Furthermore, increased chromatin condensation may indicate an impaired efficency of DNA repair activity. It has been previously described that chromatin condensation participates into the activation of DNA damage response [23], however persistent condensation can interfere with the recruitment of DNA repair machinery to specific sites, resulting in the accumulation of DNA damages. Whether damages (*i.e.*, single or double strands breaks) accumulate in the DNA of aging sperm, they can affect embryo development [13], lead to pregnancy loss [14] or contribute to increased risk of post-natal diseases [12]. To verify whether DNA damages increase over time in aging mouse sperm, in addition to the standard SCSA calculation method we tried a novel data analysis approach. Briefly, sperm DNA fragmentation includes both single and double strand breaks [24–26]. Since the intensive DNA denaturation is executed in pH=1.5 during SCSA, we assumed that more single strand fragments can be expected from DNA double strand breaks (dsbDNA) than from single strand breaks (ssbDNA). Our data showed a weak (but not statistically significant) correlation between age and increasing dsbDNA in mouse spermatozoa.

In the second part of our study, we looked at “qualitative” parameters in aging mouse spermatozoa. Our data did not show any significant effect of aging on metabolic activity (by CellROX Green assay) and apoptosis (by AnnexinV/FITC assay), as previously reported in outbreed CF1 mice [9]. Differently, other studies described age-related negative changes in human sperm quality [27,28]. One of the reasons behind this discrepancy may rely not only in the differences between species but also on the absence of health issues and exposure to environmental factors (i.e. toxin, pollution) in our model. The animal used within our study have been maintained under controlled conditions and, in case of health problems, animals have been excluded. Moreover, we observed that exposure of mouse sperm to different temperature during incubation (37□/30minutes - CellROX green assay; on ice/15 minutes – AnnexinV/FITC assay) results in percentage of intact membrane that do not overlap for individual samples, suggesting high susceptibility to environmental conditions. Thus, as men are more likely to be exposed to potentially harmful environmental factors (*i.e.*, toxins, heavy metals, pollution), the poor sperm characteristic in aged men may be the results not only of aging but also of other risk factors (both clinical and environmental).

In summary, our study showed that aging in a mouse model without underlying infertility (or other health conditions) did not affect DNA integrity and qualitative semen parameters while is associated with increasing sperm chromatin condensation. Further investigation must focus on targeted and quantity parameters (i.e., targeted DNA methylation) to better clarify whether or not aging is responsible of poor reproductive outcome in mammals.

## CRediT authorship contribution statement

**F Zacchini:** Methodology, Investigation, Formal analysis, Visualization, Funding acquisition, Writing – Original Draft; **M Bochenek:** Methodology, Investigation, Formal analysis, Writing – Review and Editing; **S Bisogno:** Investigation; Writing – Review and Editing; **A Chan:** Writing – Review and Editing; **GE Ptak:** Funding acquisition, Writing – Review and Editing.

## Acknowledgments

This work was supported by the National Science Centre of Poland (GA 2015/19/D/NZ4/03696 and 2019/35/B/NZ4/03547) and the European Union’s Horizon 2020 research and innovation Programme under Marie Sklodowska-Curie Actions (GA 834621 –NeuroAPA).

## Declaration of Competing Interest

Authors declare no conflict of interests.

## Notes

### Competing Interest Statement

The authors have declared no competing interest.

